# Biophysically motivated regulatory network inference: progress and prospects

**DOI:** 10.1101/051847

**Authors:** Tarmo Äijö, Richard Bonneau

**Affiliations:** Simons Foundation, Center for Computational Biology, New York, NY 10010; New York University, Departments of Computer Science & Biology, New York, NY 10003

## Abstract

Via a confluence of genomic technology and computational developments the possibility of network inference methods that automatically learn large comprehensive models of cellular regulation is closer than ever. This perspective will focus on enumerating the elements of computational strategies that, when coupled to appropriate experimental designs, can lead to accurate large-scale models of chromatin-state and transcriptional regulatory structure and dynamics. We highlight four research questions that require further investigation in order to make progress in network inference: using overall constraints on network structure like sparsity, use of informative priors and data integration to constrain individual model parameters, estimation of latent regulatory factor activity under varying cell conditions, and new methods for learning and modeling regulatory factor interactions. We conclude that methods combining advances in these four categories of required effort with new genomic technologies will result in biophysically motivated dynamic genome-wide regulatory network models for several of the best studied organisms and cell types.

## Large cellular regulatory networks exist, are important, and won’t go away if we ignore them

Large cellular regulatory networks, composed of thousands of protein and RNA components interacting with the chromosome to regulate transcription, are essential to coordinating nearly all cell functions [1]. All regulatory factors are in turn regulated by the same processes they comprise and under the simultaneous control of other regulation-relevant cell processes such as localization and post-transcriptional and post-translational modifications. The end result is a complex and large network of interactions that govern cell decision-making critical to all cell functions, from cell division to differentiation to response to exogenous environmental factors. Evolutionary, genomic and genetic work supports the (functional) importance of the complexity of these regulatory networks. With all of this in mind it is reasonable to state that regulatory network models can only be reduced in scale and simplified so much before they break and become incoherent, and that breaking point, once reached, still leaves us with regulatory networks of daunting complexity. In spite of the many significant challenges to working on such large systems, the fields of genomics and systems biology have developed powerful genome-wide measurement technologies that when combined with new computational methods for learning networks produce large regulatory network models that vary in accuracy, comprehension, and model complexity tremendously (depending on the biological system, technologies used, and methods used for network modeling and learning) [2] [3] [4]. This perspective piece will not provide a comprehensive review of all of these methods (or all or the large scale genomics efforts that produce the required data-sets); instead we aim to provide an overview of a few core features that the network inference methods of the future will need to incorporate.

## Large-scale network inference is the best way to approach learning new regulatory interactions even if you are primarily interested in one or a few cell processes or regulatory sub-networks

In this work network inference is an umbrella term for computational methods that identify (quantitative) gene regulatory networks (sets of regulatory interactions between transcription factors (TFs)) and genes) from experimental data. There are a number of reasons for approaching cellular regulation using a large-scale computational network inference framework. Given the scale of genomes and the correspondingly large scale and complexity of robust regulatory apparatus controlling genome structure and gene expression, learning regulatory interactions one or a few at a time is woefully inefficient and is prone to observation biases. The inherent very-large scale of functional regulatory interaction networks is shown by: 1) recent high-throughput experiments aimed at TFs (in most cases well known TFs have many more regulatory targets than known prior to genomic interrogation) [5] [6] [7] [8] [9], 2) recent high-throughput experiments aimed at determining functional regions and structure of the genome [10–14] [15–18] [10, 19–23], 3) genetics and evolutionary considerations which show that expansion and neo-functionalization of regulatory gene families is ongoing and under strong selection [24] [25] and lastly 4) by single-TF and traditional molecular biology (by continually finding and characterizing new regulatory interactions we see that the end of the regulatory complexity connected to phenotype is no where in sight).

Scale and efficiency based-arguments (akin to the above considerations) are, perhaps, the weaker and more obvious reasons to employ large-scale regulatory network inference and genomic approaches. Large-scale network inference approaches also have several advantages with regard to experimental design: that is, we can learn things from large-scale experimental designs that we cannot from single-gene or single-pathway experimental designs. Often smaller scale (single process focused) molecular biology experiments must be selected from a very large manifold of potential experiments. This selection process is driven by biological intuition and the expert knowledge of molecular biologists. With even the keenest group of intelligent biologists this process introduces unacceptable observation bias and inhibits scientific reproducibility. The important process of selecting what to test in the first place is often poorly described and key negative results leading to this selection are omitted or buried in poorly written supplemental materials. Avoiding the observation bias of small-scale molecular biology workflows is a main attraction of genomics-powered network inference. Unbiased approaches to network inference enable cross-comparisons that are not always possible with single-gene or single-process experimental designs and better enable comparison of newly learned interactions to known interactions (and enable in some case a more unbiased reevaluation of error in our ‘known’ set of regulatory characterized interactions).

Lastly, regulatory network inference does not obviate detailed study of regulatory interactions, it enhances such endeavors, by providing a weighted map that a molecular biologist can use to select and prioritize factors to further characterize. With sufficiently detailed representations (i.e. sufficiently biologically motivated core representations of the core components of the regulatory network) we also provide opportunities for biochemical and molecular biology follow-up studies that would not exist with a single-gene approach. A main reason for pursuing methods that result in biophysically motivated models of regulatory networks is to better integrate initial network inference with downstream functional characterization of the bigger better regulatory interactions one catches with network inference.

## Method diversity in network inference stems, necessarily, from the diversity in how we model regulatory networks and the diversity of genomic technologies and experimental designs

There is no one-size-fits-all solution to network inference [26] [27–33]. There are a large number of methods for network inference due to the importance of the problem and the diversity of experimental designs. For example, the experimental designs and computational methods available to scientists studying bacteria without good genetic systems differs from methods appropriate for scientists studying the immune system in human. The appropriate method for network inference must be determined after a careful consideration of several factors including: 1) the biological questions that are priorities for any given study (do you want to predict the effects of knock outs on metabolism or find new regulators with a predefined phenotype?), 2) the orthogonal measurement techniques available and applicable, 3) the budget and the scale of the project, 4) the availability and diversity of public data, and 5) the extent of prior knowledge about the system. These considerations affect the model simplifications and requirements that are allowed/needed and thus should influence method development and selection. Computational modeling should not only be used as a post measurement analysis step. Instead we should carefully match experimental design to the computational tools we plan to use if a major motivation is optimal discovery of new regulatory interactions with large-scale (unbiased data-driven) methods.

## The scope of this perspective

This perspective does not intend to provide a broad and comprehensive review of methods for network inference, but instead will express a very focused view on where the field of regulatory network inference needs to go in order to build genome-wide biophysically motivated regulatory network models. We also aim to provide the reader thoughts on how to coordinate experimental design with computational methods for network inference. In this section we will explicitly state our assumptions, restrictions and focus. For a more comprehensive discussion see any of these recent reviews of network inference in biology. [27, 32, 34–44]

Our first and largest restriction of focus is that we will exclusively discuss regulatory networks controlling transcription and chromatin state. We will consider methods that learn networks of factors that regulate chromatin state, chromatin structure, rate of transcription at target genes, transcription factor activity and RNA-degradation and processing rates. That is to say, that our models stop at translation (precisely where technologies beyond nucleotide sequencing are needed to make measurements). Networks of protein interactions and the regulation of translation and post-translational processes are clearly an exciting and an important area of research and also amenable to large-scale computational methods for discovering and characterizing interactions. Modeling of metabolism and its interaction with gene-regulation is also omitted here. We feel this scope is justified as: 1) several of the discussions here can be in part applied to other informational levels given appropriate data and 2) transcriptional network inference together with progress in mapping regulatory chromatin will aid ongoing efforts to integrate models across these other informational levels. We also believe that no single approach is likely to be optimal given the distinctly different properties of different types of biological networks. Given this focus, we will discuss recent complimentary genomic technologies that are driving the next generation of integrated experimental designs and computational approaches to transcription and chromatin-state regulatory network inference.

We will focus here on network inference methods such as Inferelator that result in models with units of time and concentration (or relative concentration). Transcription and its regulation are dynamic processes by their core nature and any approach to modeling gene regulation that does not take dynamics (or at least timing) into account is, in our opinion, leaving out key biology. Uncertainty about the critical timing and rate of events here motivates the need for time series and dynamical experimental designs as uncertainty about the interactions motivates genetics and perturbation driven genomic approaches to discovering regulatory interactions.

## Defining network models and determining model complexity

In many fields of computational biology, the main factor separating schools of modeling/analysis is resolution. For example, in modeling protein structure, early computational work was separated into schools of thought partitioned along an axis that spanned atomic resolution molecular dynamics methods to statistical lower resolution methods[45]. As the availability of structure data, the power of computers, and the sophistication of methods progressed, lower resolution methods have been largely supplanted by high-resolution (atomistic and even hybrid quantum mechanical) methods. Although the problems of structure prediction and network inference are fundamentally different, we believe that there are some parallels to the evolution of methodologies in both fields and that resolution/detail of resulting models is a primary axis to consider. We believe that regulatory network models are demonstrably moving towards more detailed representations commensurate with the data availability and richness (and commensurate with better overall experimental designs).

### Undirected vs. directed

A first level of approximation to consider in network inference is whether the resulting network will be undirected or directed. Directed graphs are desirable in regulatory biology, as we know the regulators involved in many case, we wish to use these graphs to predict outcomes of perturbed systems. Directed graphs imply causality even when methods that do not explicitly consider causality are employed, and thus there are several reason that inferring directed graphs is markedly more difficult than undirected graphs. Although several notable examples of the use and learning of undirected graphs exist this review is written with the overriding idea that directed network representations, although difficult to get right, are the eventual or immediate goal of regulatory network inference. The idea that directed graphs are needed is driven by the assumption that regulatory models should be used in forward engineering and genetic-interpretation applications (where directionality and ideally causation are desired model properties).

### Discrete-vs. continuous-valued

There are several advantages to reducing the representation of gene-levels to discrete (1 and 0 in most cases) levels. Methods for determining steady-state behavior and fitting Boolean models to data can be quite efficient [30, 46]. Current methods based on (dynamic) Boolean network models have been shown to powerful in modeling interactions and asynchronous dynamics across networks [47, 48]. The largest advantage of methods based on these models is that the extreme representational efficiency allows for enumeration and evaluation of large (and in some cases exhaustive) sets of proper interaction terms (AND and OR interactions between TFs for example). Methods based on decision trees and forests of decision trees represent a model representation somewhat between Boolean and real-valued data representations (where there is a discrete set of values for each gene, but they are fit in the original data-space) [49]. A major potential disadvantage of these methods is that they require mapping data from continuous values (where real-valued variance is often biologically relevant) to discrete labels via a lossy transformation. This review will focus on real-valued network inference methods where the real values representing genes are assumed to correspond to rates (for model parameters), concentrations (for targets) and concentrations or activities (for regulators, where activity is given in units of concentration or relative concentration).

### Dynamic models of regulation

Many methods for regulatory network inference represent TF-target interactions as TF effect of target rates of transcription. Modeling transcription and transcriptional regulation as a convolution of many thousands of rates and influences on rates turns network inference into a large-scale parameter estimation task that at first glance seems impossible to solve. Methods for model selection, influencing model selection with informative parameters, integration of multiple-data types and proper experimental design can (we argue throughout this paper) be combined to make this problem tractable. The most commonly used differential equation models are ordinary differential equation (ODE) and stochastic differential equation (SDE) models where TF effect on transcription rate is learned using a greedy search (one target gene at a time). Several challenges remain for learning dynamic models including: 1) treating the parameterization of these large networks as a proper global system by simultaneously fitting all parameters [50]2) modeling latent states like transcription factor activity [51] [52], 3) explicitly modeling activator, repressor[53], degradation and target expression with distinct biophysically correct distributions, and 4) determining correct methods for using these models to design optimal experiments.

### Interactions

Learning introductory molecular biology often mixes the memorization of biology’s many moving parts with learning about the experiments and analysis needed to study cells. From this experience we get examples of many small network models that represent regulatory programs of combinatorial regulation involving TF-TF interactions and TF complexes. Even though we are taught about the key relevance of these multiple factor regulatory interactions we find that a few automatic methods for network inference include interactions (with a few notable/influential exceptions). There are many mathematical approaches to modeling regulatory factors that must interact to be functional (or non-functional) including 1) use of linearly interpolated logic gates using min() and max() and 2) quadratic terms, and proper biophysical modeling of complex assembly integrated into network inference core models via approximate functions (S-systems, etc.). This is due to the fact that interaction terms dramatically increase the rate at which complexity of models grows with number of predictors. In many cases current data sets do not support the de novo learning of interactions and we posit below that interactions should be included and modeled using integration of multiple data types. Network inference is typically driven by data sets that give us windows in to system state and dynamics (like expression, proteomics and phosphoproteomics) but to include interactions and properly capture the biology we must integrate interaction-centric data types (protein-protein interactions, genetic interactions, chromatin accessibility and cooperative TF binding).

When deciding the resolution of our network model (and thus deciding between the large number of potential representations and associated methods) we must balance the biological/modeling demands to be placed on the model (how we hope to use the model) with the constraints on what we can learn imposed by technology, the scope of the study and the experimental design. Learning more detailed models can increase the need for more elaborate experimental designs at larger scale resulting in increased cost and work levels. In many cases biological systems are not amenable to repeated interrogations or genetic manipulation, which confounds the inference of more detailed and more comprehensive networks. Thus, part of the motivation to use simplified representation stems from the need to learn networks from small data sets and from imperfect experimental designs. We must not err on the side of oversimplifying our models, as this will limit proper comparisons to data and the usefulness of the model revealing novel biology. Recent works show that network inference tasks are not inherently harder than undirected graph identification tasks. This notion is supported by the success of dynamical models in blind prediction settings [29] and in the context of inference of networks that were then experimental validated via large-scale functional follow-up experiments [5] [7].

There are several ways in which increasing the detail and biophysical relevance of our model can improve or enable accurate learning when compared to less detailed models. More detailed representations of regulation can, however, have several advantages with respect to learning from limited data sets. The most direct advantage is that more detailed representations may require fewer assumptions to compare model states to data (and thus fewer lossy data transformations with inbuilt assumptions are required). An example would be comparing a continuous dynamic model to a Boolean representation of transcript levels. In the case of the Boolean model the parameter space might be significantly smaller, but severe lossy assumptions are needed to transform the original observations into the binary space of the model. Another advantage is that additional model complexity (especially with respect to dynamics) can allow for more direct integration of data from different studies and different measurement technologies.

More detailed representations can also have several advantages when integrating network inference methods into the larger experimental & biological context of a large-scale study. For example, including modeling of dynamics and time might allow for additional flexibility in designing time series experiments (where other methods might perform best with regularly sampled time series), capturing important dynamics from irregularly sampled time series and might allow for prediction of data for time series and other experiments that have significant missing data (afforded by explicit modeling of time and transcriptional dynamics).

## Relevant genomics technological advancements and biological data

Before going any further, we focus on a minimal set of data types needed to genome-wide chromatin & transcriptional network inference. These data types fall into four broad categories: expression data, chromatin accessibility data, chromatin structure data and TF-DNA interaction data. Expression data can be used to derive core regulatory models while the former three data types provide information on TF-loci interactions that can be used to select correct network models and estimate TF-target activity relationships.

### Expression/state

Gene expression, a key staple data type for regulatory network inference, is also the state variable for the target genes in many of the models we describe herein. For transcriptional regulatory network inference these state variables are measured primarily by microarray and RNA-seq, which aim to make comprehensive genome-wide measurements of different RNA molecule types. Other key technologies are emerging that promise to allow direct measurement of RNA-degradation [54], measurement of actively transcribing transcripts, integration of transcription and proteomic measurements [55] [56, 57] and other more kinetically relevant views on transcription. As our focus here is on chromatin and transcription networks we leave out any discussion of the intriguing current work to connect transcription and translation [58] [59]. These technologies can be combined to form multi-view data sets, that optimally combine perturbation, genetics and time series experimental designs into an overarching design. Overarching experimental design considerations, in the context of network inference[60], are discussed more below.

### TF-binding and chromatin state

Key to tackling the scale of regulatory networks are means of identifying active TF-binding sites from putative binding sites in the genome and ways of reducing the set of all genes to the set of active (or differentially regulated) targets in any given cell or cell-state. Key data for this model-space reduction are data sets that tell us where TFs bind the genome. Several such experiments (ChIP-seq, ChIP-chip [61, 62], ChIP-exo [63, 64], ChIA-PET [65] and others) exist and can be combined with expression data to derive detailed priors on the structure of the network. A key limitation is that these experiments target a single TF at a time and thus do not scale easily when hundreds to thousands of TFs might be involved in any given process of interest. Imperfect, but very highly scalable, alternatives can be found in methods that combine the TF-binding motifs [66–69] of TFs with chromatin state [70] (including histone modifications and chromosome accessibility, where accessibility = nucleosome free and possibly bound by regulatory factors). Motifs found for a given TF in an accessible region are much more likely (although far from guaranteed) to be active or functional. The availability of >1000 motifs for mouse or for human, for example, allow us to extract from each accessibility experiment the network-inference equivalent of 1000+ noisy ChIP-seq experiments [66–69] [71–73]. Key limitations include the fact that many homologous TFs share nearly identical binding sites, and like with ChIP-seq, a small fraction of binding sites are thought to be regulatory in the cell’s current state. Thus, to scale the estimation of structured informative prior we need new computational methods that can deal with this error and large-TF-family ambivalence.

### Chromatin structure

Another problem in regulatory network inference is the assignment of binding sites and regulatory genomic loci to their targets throughout the genome (actual sites of transcription). In many cases methods applied to large integrated data sets make the assumption that binding sites near genes are potentially involved in regulating proximal genes (throwing out all interactions that skip over genes and throwing out all long range interactions along the chromosome). This limiting assumption was, prior, needed to limit the number of putative interactions and preserve the main intended use of data like ChIP-seq (that the data reduce the space of plausible models). New data sources allow us to estimate chromosomal structure through both the explicit measurement of interactions and through the identification of chromosome neighborhoods along the genome [74–76] [77, 78]. With the increase in the quality and prevalence of these data sources we will find that weaving these data types into integrated experimental designs will offer the ability to connect enhancers to distal regions accurately and without the model size explosion.

## Example experimental designs for networks inference

When approaching how to design experiments for (or curate inputs to) network inference we have to consider both the balance between the experimental types and the possibility that several large efforts may have produced relevant data sets already. Experimental designs for network inference need to balance all of these data types, but how to construct optimal experimental designs [79, 80], given a network inference method and such diverse TF-binding, accessibility and chromosome-structure data types remains an open question [81, 82]. Biological constraints also strongly restrict experimental design in unexpected ways; for instance, we want to see what that gene knockout (KO) does to T-cells, but knocking out that gene is developmentally lethal. Two recent experimental designs for network inference are presented below (both examples where binding/constraints are balanced with state/expression data). In both cases we do not imply that these designs are examples of optimality: both of these studies enabled network inference but were not designed solely for network inference. There were several time and reagent cost constraints; and we have learned a lot about the computational tools in the last five years that would lead to improvements in these experimental designs. That said, these studies are the result of balancing network inference experimental design against practical and biological constraints with the aim of learning large-scale regulatory network models.

### An example experimental design for studying T-cell differentiation and function

In our recent work aimed at learning T-cell networks that coordinate and drive T helper 17 (Th17) differentiation and function we combined time series and KO RNA-seq data, and ChIP-seq data types to constrain a plausible regulatory network model [5] [83–86]. This resulted in identification of several novel TF-target relationships important to the immune-phenotypes we were interested in. Approximately 1/4 of >2000 identified regulatory interactions were tested and validated by knocking out TFs or performing follow-up ChIP-seq on TFs found with the model. The core of this data set was a set of >150 RNA-seq experiments aimed at different T helper (Th) cells (e.g. Th0, Th1, Th2, and Th17) and T regulatory cells (Tregs) with the majority of the data collected for Th17 cells. This data was split nearly evenly between a time series following Th17 differentiation, KO of ∼30 TFs, and several other key perturbations. Added to this were ChIP-seq experiments performed for ∼30 TFs (mostly in house). Lastly large numbers of public data sets were used, including a large atlas of gene expression in other immune cell types (the ImmGen data set, as well as several recent single-cell RNA-seq, knockout and orthogonal RNA-seq experiments could be integrated into this data set) [3, 87–90] [91, 92]. As we have developed new computational and genomic techniques, the Littman lab has now (post the 2012 work) added to this 35 ATAC-seq (an assay for transposase-accessible chromatin using sequencing) experiments for mapping active chromatin in different cell states, hundreds of additional expression experiments in a broad array of cell types and single-cell RNA-seq experiments (Dan Littman, unpublished). Thus, sufficient data exists for the estimation of hundreds of TFs and chromatin remodeling agents in hundreds of lymphoid cell types and states (using ChIP-and ATAC-seq to derive noisy priors on network structure and then using those priors to estimate TF activity, see below). The expression data, when combined with informative priors derived from binding and accessibility experiments, enables inference of detailed and dynamic regulatory networks. A main missing element in the T-cell network data-corpus is information about chromosomal structure that can be used to bring distal enhancers into regulatory network models, but current efforts are rapidly closing this gap.

### An example experimental design for B. subtilis

Another example comes from the world of bacterial systems biology. Two studies aimed at large-scale regulatory network inference resulted in huge strides towards a global regulatory network model of the *B. subtilis* (thousands of newly discovered regulatory interactions) [7, 93, 94] [8] [95, 96]. In this organism ∼2,900 regulatory interactions were known prior to these studies, and so both attempts could rely on large numbers of ‘known’ interactions for tasks like generating informative priors and estimating TF activities. The use of such a large number of ‘known’ edges to generate constraints on network structure and to estimate TF activities is in strong contrast to the above experimental design (where ChIP-seq and ATAC-seq were used to generate network priors and ‘known’ interactions were kept as a mini-benchmark). Databases like RegulonDB [97] for *E. coli* and SubtiWiki [98] for *B. subtilis* containing experimentally verified interactions are extremely valuable resources for both developing and applying network inference methods in these and closely related organisms. It is important to note and emphasize that some of the earliest and most successful studies aimed at global models of regulation and regulation’s interaction with other systems in the cell were carried out in bacteria and archaea [8] [6] [9] [96]. The main body of this data set consist of two sets of microarray experiments (one on a tiling array, the other a custom Agilent array). In both cases the arrays were able to also detect long noncoding RNAs as well as mRNAs. One set of experiments focused on a set of perturbations followed by sampling of short time series, the other was a mix of genetic and metabolic perturbations (although both datasets were very diverse in the shear number of environmental and genetic perturbations explored). From combining these data sets (as well as analysis on subsets of the data) multiple computational groups were able to expand the global regulatory network, estimate activities for approximately half of the TFs over all the conditions explored, and even learn dynamic regulatory network models.

A key distinction between this work in *B. subtilis* and prior work in *Halobacterium* [6] is that genetic perturbations in *Halobacterium* (an archaea model system) (as well as several current genomic technologies) were not available prior and the number of known interactions were quite small. Therefore, the use of genetics and informative priors was not possible in 2005, resulting in a distinctly different time series and environmental perturbation based approach aimed at accurately predicting trends in data (such as future time points in time series). This contrast highlights nicely the degree to which biology ultimately (and correctly) dominates experimental design except in cases like human or model systems where vast human efforts have developed genetics and other systems needed to dissect these systems at will. Similar contrasts exist when considering experimental designs aimed at learning and comparing human and mouse regulatory programs. Model systems are still (and might always be) key to progress in this and other fields of biology.

## Our four part recipe for network inference

For the rest of this perspective we will present our four part recipe for breaking the problem of network inference into approachable sub-problems and working methods given the data we can reasonably expect to be able to collect and integrate with public data. The four sub-problems are: 1) using structural assumptions like sparsity that effect global network properties, 2) generating and using informative priors on network structure that effect single edges, 3) TF activity estimation, and 4) using guided/intelligent TF-TF interaction terms.

## Model selection in network inference using assumptions about sparsity, structure, rank etc

Given the significant challenges we face when learning regulatory networks from biological data sets of limited size and constrained design we must develop computational methods that take full advantage of both biologically motivated assumptions and data integration in equal measure to solve the problem[99]. One of the key assumption is that biological networks are sparse [100]; that is, to say that most regulatory factors do not regulate most genes, thus the true regulatory network has many fewer interactions than the full set of possible interactions. This constrain has been built in one way (Lasso, various information criteria and marginal likelihood etc.) or another into nearly every method currently in use. Key places to look for discussions of sparsity constraints in model selection in biology include: [101] [102] [103, 104] [105, 106]. Given that this a mature area of discourse and part of nearly every method cited here we will not discuss or compare the myriad methods.

In spite of our best efforts to identify networks with simple topology priors, like sparsity, there are several aspects of biological networks that confound our efforts. One major problem is that several sets of genes are co-expressed [107]. Thus, a TF and a potential target gene may have significant correlation, mutual information or other similarity metric, due only to the fact that they are co-expressed (thus several false positives could result from misattributing co-expression as direct regulation). Two possible classes of approaches to this are pre-clustering (or pre-biclustering) [94, 108–116] and the use of time series and dynamics to differentiate co-regulation from true regulatory interactions (as co-expressed should appear instantaneous and true regulatory interaction should incorporate a delay between TF activity and effect on target) [117]. These cases would include fan-out motifs where multiple of the downstream targets are in turn TFs.

Other motifs including feed-forward, feed-back and auto-regulatory loops also present considerable difficulties. In recent double blind tests of network inference methods several such loops were identified as especially difficult to infer correctly across all methods tested [29, 118]. This is consistent with several previous works that show that even small signaling and regulatory networks can suffer from non-identifiability (e.g. where the true network generates data consistent with a large number of model structures). A last consideration is that biological networks are in many cases robust to perturbations including deleting nodes and edges and modifying strength of edges. This biological robustness to perturbations implies that removing components of the true network will result in incorrect models that are equally consistent with the data when a few observational methods are used [119]. Although difficult to prove degrees, this line of thinking suggests that many learned regulatory network models are in fact overly sparse. With all of this in mind, we can safely state that constraining network complexity, although an important component of most methods, is only part of the recipe to the network inference problem.

## Model selection with informative priors: using constraints to fight the non-identifiability problem

Given the shortfalls of data sets, unavoidable biological constraints and the likely non-identifiability problem discussed above we must turn to orthogonal data sources and prior information to aid in selecting correct models from the large number of candidate models consistent with our data. Prior information about network structure can come from a huge variety of sources. First and foremost are ‘known’ interactions; in *S. cerevisiae*, *E. coli* and *B. subtilis* we could use thousands of previously characterized regulatory interactions to influence our network inference [120] [121, 122] [97, 123] [98, 124–126]. We also look to experiments like ChIP-seq and others described above to provide information about where factors bind in the genome. With these experiments we still have several primary analyses to perform before suitable network priors/constraints are derived. These primary analyses involve matching peaks to target genes hampered by lossy assumptions including ignoring or incorrectly mapping distal influences. More recent, computational methods combine chromatin accessibility experiments with regulatory factor DNA binding motifs to derive priors on connectivity for all TFs with known motifs simultaneously. Thus, the prevalence, accuracy and coverage of structure priors and methods for deriving informative priors on network structure is rapidly increasing.

Computational methods that aim to use data integration and known interactions to constrain network inference must face a high error/irrelevance rate (many binding sites are not functional or they are functional in other cell states/types) and must deal with heterogeneity in the evidence for individual edges (the high variability in ‘known’ edges).

Recent work in our group used informative priors to modify both shrinkage based methods (modified elastic net) and Bayesian linear regression with modified informative priors on network structure (Inferelator-BBSR) [7, 127] [128]; in both cases we were able to show that our methods improved our ability to infer correct network edges even when signal to noise ratio was as low as 1/10 (where signal to noise ratio = number of regulatory relevant edges divided by relevant edges + noise and non-functional). Several other groups have also recently made progress in using informative network priors both within computational biology and statistics[129] [130–132] [133]. It is clear that informative and structured network prior generation and use will remain a fruitful research topic in the future [129] [130–132] [133].

Using new high-dimensional statistics methods to couple related inference tasks (such as inference for closely related species, closely related cell types, and similar cells interrogated with different technologies) also promises to provide new ways of identifying correct network models [134–136]. Several approaches have been proposed to coupled model identification and fitting that incorporate inter-model distance to the overall penalty (objective) function. These inter-model distances reward orthologous pairs of TF-target interactions that have similar parameters across models. A simple approach to this problem for multiple closely related species would be to select TF-target pairs believed to be conserved and add a penalty term that pushes coupled (conserved) model parameters toward similar evaluation (some function of the difference between the model parameter in mouse and human for example) [137, 138]. However, coupling inference tasks runs the risk of forcing interactions that are no longer conserved to manifest as false positives or false negative in either task. Newer versions of these methods need to be developed that allow for this constraint to be learned in ways that allow for orthology constraints and other linkages between inference tasks to be down-weighted when not supported by the data[139]. For instance, when TF-target pairs have been neo-functionalized or when regulatory interactions do not span the cellular conditions represented by distinct inference tasks [140] [141].

## Estimation of transcription factor activities: using data integration to turn every cell into a massively scaled reported assay

Genomic technologies advance at a tremendous pace; a key misconception this innovation fosters is that a technology that can measure a desired cell property is right around the corner. This misconception limits the degree to which groups embrace proper integrative multi-data-type experimental designs and analysis frameworks. A key example comes when we consider that transcription factor activity is perhaps one of the most important variables in the cell (especially if we are interested in differentiation and adaptation). The activity of a TF (or any protein or functional RNA) is the concentration or active molecules (or relative concentration) [142]. For STAT3 in mouse this would be the concentration (presumably a fraction of the total protein between 0 and the total concentration of STAT3 protein) of STAT3 that is phosphorylated and in the nuclease [143]. For RORC in mouse this would be the concentration or this nuclear hormone receptor that has ligand bound and is located in the nucleus [143]. Many other transcription factors need to interact with other proteins to form active complexes (e.g. IRF4 and BATF in mouse) [5] [144]. It is important to remember that activity also depends on the state of the chromatin target loci [145, 146]. This activity (in these diverse cases) can be represented with the same diversity of representation that we use for TFs and target expression: from Boolean all the way to concentration proper with all sorts of normalized relative representations in between. Looking at a few examples we find that the activity of transcription factors cannot be simultaneously measured directly in a single uniform assay or set of assays for all transcription factors.

The key to efficiently estimating transcription factor activities genome-wide is data integration [7] [147–159]. Consider how we would estimate TF activities if we had a perfect regulatory model: we would simply plug in the measured expression observations and solve for the TF activities. Strikingly, this would allow us to utilize known regulatory interactions to turn each set of RNA-seq experiments into massively parallel reporter assays reporting on TF activities. In practice, we are faced with estimating TF activities from noisy data with a very incomplete set of regulatory interactions. There are several groups that have demonstrated progress in developing methods, that with real data, can derive activity estimates that match assays of activity for select TFs[7] [147–159]. Importantly, these estimated TF activities improve downstream analysis and ultimately improve the accuracy and coverage of regulatory network inference. Early methods utilized linear regression approaches to solve this problem. More recent methods range from detailed probabilistic methods that can take advantage of the dynamics captured in irregular time series data to the use of earlier methods embedded in more recent network inference and model selection methods. We believe that explicit estimation of the activity of regulatory factors must be a central part of any future successful network inference method, but several open questions remain. Current approaches include methods that first estimate activities and then learn networks based on those activities. We believe that future methods need to integrate network and regulatory factor activity estimations. Another key question to consider (at least in eukaryotes) is the receptiveness/accessibility of the target loci and the possible need to model this target-receptiveness when estimating TF activities.

## Need for interactions: cell type specificity of regulatory networks is overemphasized in current works

A key question in systems biology is: to what degree do we need cell-type and cell-state specific network models. Here we posit that the level of representation used to model regulatory networks is a key to answering this question, and that more detailed biophysical models can span larger numbers of cell types than less detailed representations. An extreme example would be a ‘perfect’ model of a mouse embryo or a bacterial cell. With this model in hand we could generate any cell type or state and then simulate its behavior, thus we would be in possession of a model that spanned all cell types. On the other end of the cell-specificity-need spectrum we find current practice, where labs collect large data sets with the aim of making network models relevant in small numbers of cell states (for example learning separate models for different finely divided types of T-cells). In practice, we find that many labs pursuing single celled organisms pursue global models while multi-cellular organisms are often dominated by efforts aimed at ever more distinct cell sub-types (presumably leading to equal level of complexity). Like all questions posed in this perspective, there is no one-size-fits-all solution, however we believe that increasing the detail and biophysical motivation behind models would decrease the need for different models for each cell type[160].

Modeling interactions between regulators and chromatin state is perhaps the most important level of detail needed in current models (not found in most regulatory network models to date) because it is a prerequisite for formulating multi-cell state models[146, 161, 162]. There are a large number of challenges that have limited the degree to which true interaction terms appear in network inference models. This is due to both the need to specify the type of interaction (AND, OR, fuzzy logic like ?ukasiewicz logic, etc.) as well as deal with the large explosion in the possible model complexity that allowing the number of predictors squared terms into a model. Typically, interaction terms have been included in models that have reduced model complexity, such as Boolean models and asynchronous Boolean models in which huge gains in representational and model-evaluation efficiency allow for searches over very large spaces of possible interactions. Early versions of the Inferelator included such interaction terms for environmental and TF-TF interaction terms (linearly interpolated AND gates) [163]. In addition to improving cell-type comparability, proper interaction terms should also increase the degree to which the network representation matches known key biological interactions that are essential to several of the best characterized regulatory networks.

## Frameworks for putting it all together

How do we build a method that encompasses structural constraints, informative priors, TF-TF interactions, and TF activity estimation? How do we put the pieces together? One might respond to either of these questions in a number of ways, but for simplicity here we will reduce this overall field-wide question to a simpler variant: what is the value in simultaneously estimating these qualities as part of a network inference procedure? We might break the calculation up into modules, whereby activity estimation, TF-TF interaction selection, and network inference are carried out in three separate modules [7], or we might construct a procedure that attempts to solve these estimation tasks simultaneously within a single unified model.

In principle, gene transcription and its regulation could be modeled rather accurately with exact stochastic simulation of coupled chemical reactions [164, 165]. Unfortunately, the practical usefulness of the this approach is severely impaired by many practical facts, e.g.: 1) it requires a detailed specification of the chemical reaction network model (the details are not known), 2) model inference requires single-cell data on many molecules (not feasible in practice), or otherwise the model is rendered by many latent variables, 3) many reaction parameters are unknown and difficult to measure, 4) inference of a large-scale model is computationally intractable and 5) model identifiability is questionable due to the flexibility of the detailed model rendering model selection difficult. For these reasons, ODEs and SDEs have been popular for modeling gene transcription on a population level and its regulation through transcription factors, and for inferring cell-type specific transcription factor networks in the data-driven manner[36, 166, 167]. Undoubtedly, the TF focused view of transcriptional regulation is a crude simplification of the biophysical phenomenon, but despite that, it has been empirically shown to be efficient and yet accurate approximation. Additionally, the importance of TFs in regulating transcription is undisputed, and their mechanisms of action on transcription regulation are well understood [1, 168].

Implicitly, many (among our) TF centric approaches are based on the assumption that TFs collectively modulate RNA polymerase II (RNAPII) recruitment and elongation (could be measured using ChIP) resulting in changes in rate of gene transcription [51, 169, 170]. Importantly, traditional mature mRNA sequencing (RNA-seq) results only in snapshots of total mature mRNA levels, that is, the rates of transcription are not directly measured (could be measured using GRO-seq/ chromatin associated RNA-seq). Hence, many existing statistical models consider rates of transcription as latent variables affecting total mature mRNA levels together with basal/degradation rates[171–173]. Finally, in a biophysically motivated model, the rate of transcription should be strictly non-negative, and thus the loss of mature mRNA is explained solely by the degradation (accessible by inference and recent experimental techniques).

Many current methodologies explicitly assume that the effect of a TF on transcription is a function of concentration of its active form and protein activities have to be considered as latent variables because (as discussed above) of the difficulties in measuring activity directly genome wide. Initially, mRNA levels of TFs were assumed to work as proxies for TF activities; however, this assumption is inaccurate in general due to nonlinearities in post-transcriptional regulation and protein translation, phosphorylation, and translocation. Importantly, this limitation can be overcome to some extent since we have demonstrated that the preprocessing step where estimation of TF activities are computationally estimated by utilizing experimentally verified regulatory interactions improves the precision of overall network inference[174, 175].

The challenge of gene transcription network inference stems significantly from the fact that the rates of transcription (measurable in principle) are convolutions of contributions of many TFs; for instance, transcription of a gene can be driven cooperatively by several promoting and suppressing TF-DNA interactions at promoter and enhancer/insulator regions (even a single TF could affect through multiple TF-DNA interactions). Identification of the target gene of a distal regulatory region is difficult because distal intergenic regulatory regions can reside megabases away from the affected gene; although, it has been demonstrated, that the analysis of enhancer and promoter transcription patterns across numerous cell types and perturbations can be used to identify target gene and enhancer pairs [176]. Alternatively, one can obtain information on the 3D organization of chromosomes in the nucleus [77, 177]; arguably, active distal regulatory regions should be located close to the target genes. Genome-wide chromatin accessibility can be used to identify potential enhancer regions; chromatin accessibility can be measured using e.g. ATAC-seq [18]. ATAC-seq has been demonstrated to be a relatively cheap and fast method for identifying temporal and cell-type changes in chromatin accessibility[178, 179]. However, ATAC-seq even when combined with motif-based sequence analysis tools does not necessarily identify which TFs are bound[180].

Generative models are statistical models for generating observable data given relationships between variables and corresponding parameters. Because generative models define hierarchical relationships between variables, they can be advantageous in integrating different data types. As always, it is important to match the level of detail of the model with the data and modeling objectives. One could formulate a hierarchical generative model for inferring TF networks by integrating temporal 5C, ChIP-seq of many TFs, and RNA-seq data. To accomplish this, the variables in the model and their types should be defined carefully; for example, 3D organization of chromosomes could be modeled using a binary matrix representing interactions between chromatin domains, TF-DNA interactions could be modeled using binary variables representing interactions between TFs and genomic regions, target gene and enhancer pairs could be modeled using binary variables representing pairs of target genes and genomic regions with bound TF(s), and TF activities and gene expressions could be modeled using continuous variables. Next, the relationships between variables within and between hierarchical levels should be defined; arguably, target gene and enhancer pairs should be compatible with the 3D organization of chromosomes, that is, interacting target genes and enhancers should reside within a chromatin domain or in interacting chromatin domains. Additionally, possible changes in target gene and enhancer pairs should be supported by gene expression and TF-DNA interactions; that is, appearing and disappearing target gene and enhancer pairs should have an effect on the expression of the target gene. The effect of an enhancer on the target gene is a function of the activities of the bound TFs. TF activities can be estimated by conditioning the model on known regulatory interactions and gene expression data (depending on the model formulation and assumptions, even novel interactions could tell us something about TF activities); resulting TF activity estimates are prone to be noisy and thus it is important to incorporate and propagate their uncertainty throughout the analysis. Finally, the TF-centric approach together with TF activity estimation and target gene and enhancer pairs will help us to learn the directionality and sign of the interactions.

It is questionable whether the simultaneous modeling of 3D chromatin structure and TF-DNA interactions is worthwhile when the increased computational complexity is taken into account. For example, uncertainty in detected TF-DNA interactions is relatively small, and thus it will not have a great effect on the final estimates. One could analyze 5C, accessibility and ChIP-seq separately and in advance of the TF network inference, and then the obtained results could be used for specifying priors for the TF network inference. For instance, 3D chromatin structure can be seen as an informative prior connecting target-gene and enhancer pairs to be used in any methods that can currently incorporate priors on network structure.

Probabilistic hierarchical generative models are naturally modular and deal properly with latent variables such as TF activities. Resulting models are flexible and model components can be added and removed based on the available data and data types. In principle, it is possible to add a hierarchical level for different cell types, which would model variation between cell types and could be used to quantify how different the transcriptional regulatory networks of different cell types are. A proper statistical inference of the model propagates the uncertainty in the variables and data through the model resulting in estimates with corresponding uncertainty. Although challenges remain in scaling such complex models to larger network inference tasks, this is an area of active research and several optimization and modeling advances will likely make these detailed probabilistic models a good choice going forward.

## Coda

Several challenges remain to determining optimal overall experimental designs and computational methods for learning large-scale regulatory networks. Determining methods for integrating the estimation of disparate model components is an open area of research, but early progress in stage-wise approaches (integrating TF activity estimation as a pre-processing step) provide encouragement that such research will bear fruit and lead to improved computational methods. Another key consideration is the integration of single cell and population level genomics data. As single cell genomics becomes readily available our ability to estimate key network model components from these data will likely likely also increase, building on the network inference work carried out on populations of carefully sorted cell types and stages. New methods that integrate current datasets with single cell data sets will need to be developed to deal with the fundamentally different error structure present in current single cell data-sets (particularly missing data manifesting as extreme zero inflation) and these methods will need to co-develop with these rapidly evolving technologies. Given all of these simultaneous genomic and computational developments it is reasonable to argue that we are entering a new phase in systems biology that will see a substantive leap in the accuracy and biophysical detail of large-scale regulatory network models.

